# RAD51 nuclear recruitment and inhibition towards innovative strategies against pancreatic cancer

**DOI:** 10.1101/2021.02.03.429564

**Authors:** Fabrizio Schipani, Marcella Manerba, Roberto Marotta, Arianna Gennari, Francesco Rinaldi, Andrea Armirotti, Marinella Roberti, Giuseppina Di Stefano, Walter Rocchia, Nicola Tirelli, Stefania Girotto, Andrea Cavalli

## Abstract

RAD51, a key player in the homologous recombination (HR) mechanism, is a critical protein to preserve genomic stability. BRCA2, upon DNA damage, promotes RAD51 fibrils disassembly and its nuclear recruitment.

Here, we use BRC4, a peptide derived from the fourth BRC repeat of BRCA2; BRC4 induces RAD51 defibrillation through a ‘domino’ effect, eroding fibrils from their termini, and yielding monomeric RAD51. The congruence among several techniques (static and dynamic light scattering, negative staining transmission electron microscopy (TEM), and microscale thermophoresis) allows an accurate estimation of the kinetic and thermodynamic parameters of this process. BRC4 lacks, however, a nuclear localization sequence; therefore, it cannot transport RAD51 into the nucleus, thus behaving as a RAD51 inhibitor. Cellular assays (BxPC-3, pancreatic cancer cells) indeed show that BRC4 efficiently inhibits HR and enhances the cytotoxic effect of cisplatin, a DNA-damaging drug.

The present study sheds further light on the complexity of the HR pathway, paving the way for designing peptide and small organic molecule inhibitors of RAD51 as innovative anticancer and chemo/radiosensitizer compounds.

## Introduction

Genomic stability is key for cell replication and survival. In the cell cycle, external and internal stressors subject DNA to different insults. To address these threats, the cell has evolved mechanisms to detect DNA lesions, signal their presence, and promote their repair (Jackson & Bartek, 2009; Rouse & Jackson, 2002). The DNA damage response (DDR) pathway is the ensemble of these mechanisms. It is a highly regulated machinery that guarantees cell survival. A defect in the regulation of any of this machinery’s components often results in genomic instability, which predisposes to cancer onset (Jackson & Bartek, 2009).

Cancer cells are genetically more unstable than normal ones, and their fast proliferation strongly relies on the DDR pathway. Homologous recombination (HR) is an error-free mechanism that repairs highly toxic DNA double-strand breaks (dsDNA). HR activity has been reported to be elevated in cancer cells, where an increased rate of mutation and progressive accumulation of genetic variation over time has been observed. (Shammas *et al*, 2009). Therefore, HR inhibition may result critical for increasing DNA instability and limit cancer progression (Torgovnick & Schumacher, 2015). Moreover, because HR activity is constitutively higher in cancer cells than in normal cells, targeting the HR pathway should guarantee selectivity and reduced toxicity.

HR’s key role in fighting cancer has also emerged in connection with several anticancer chemotherapy agents and ionizing radiation specifically designed to induce DNA single- or double-strand breaks in cancer cells (Gavande *et al*, 2016). If the HR pathway performs well, cancer cells can recover from DNA alterations rather promptly, leading cancer to be resistant to radio/chemotherapy. Conversely, inhibiting HR makes cancer cells more sensitive to anticancer therapies. Thus, the inhibition of HR majorly impacts cell survival and may significantly increase sensitivity to anticancer drugs (Krajewska *et al*, 2015).

RAD51 and BRCA2 are two key HR players and are strongly associated with cancer onset and anticancer therapies. The overexpression of RAD51 has been reported in several cancers and correlates with higher and more efficient HR (Klein, 2008; Richardson *et al*, 2004). Moreover, germline mutations of the *BRCA2* gene increase susceptibility to breast, ovarian, pancreatic, and other cancer types (Petrucelli *et al*, 2010), which are more sensitive to radio/chemotherapy. Indeed, mutated BRCA2 protein can hamper the proper formation of the BRCA2-RAD51 complex and can lead to radio/chemosensitization (Falchi *et al*, 2017).

RAD51 is a recombinase of 339 amino acids that can be located in both cytosol and nucleus (West, 2003). It oligomerizes on single-strand DNA (ssDNA) searching for homologous DNA to repair damaged sequences (Haber, 2018). BRCA2 is a protein of 3418 amino acids, essential for the recruitment and accumulation of RAD51 in the nucleus repairing foci (Davies *et al*, 2001). The BRCA2-RAD51 interaction takes place through eight BRC repeats comprising 35-40 amino acids. BRC repeats are well-conserved in different species, and mutations in some of them confer sensitivity to DNA damage, suggesting their critical role for BRCA2 function.

RAD51 is mainly present in an equilibrium between monomers and homo-oligomers in the cytosol in normal cycling cells. In the presence of DNA damage, BRCA2 interacts with RAD51 and transports it into the nucleus at the site of DNA damage. This process occurs through a free nuclear pore entry mechanism or binds to other proteins containing nuclear localization sequences (NLSs) (Jeyasekharan *et al*, 2013). In the nucleus, BRCA2 assists RAD51 fibril formation along the damaged DNA, S the 3’-5’ directional growth of the fibril (Shahid *et al*, 2014). The C-terminal region of BRCA2 is also able to bind RAD51 and so stabilize RAD51 oligomers, which is needed for recombinase activity (Esashi *et al*, 2007). Once the DNA is repaired, BRCA2 disassembles RAD51 fibrils into protomers. In the absence of DNA damage, the interaction with BRCA2 prevents RAD51 oligomerization, and the C-terminal of BRCA2 is phosphorylated and inactive (Esashi *et al*., 2007). The level of nuclear RAD51 is due to regulated changes in its subcellular distribution. BRCA2 orchestrates the nuclear availability of RAD51, which enables appropriate levels of HR. By binding RAD51, BRCA2 obscures its nuclear export signal (NES), thus promoting RAD51 nuclear retention when necessary (Jeyasekharan *et al*., 2013).

The crystal structure (PDB: 1N0W) (Pellegrini *et al*, 2002), formed by RAD51 (without the first 97 residues, which are extremely flexible) linked through a peptide to the fourth BRC repeat of BRCA2 (i.e., BRC4), allowed the identification of two regions suited to the physical interaction between the protein and the peptide. The BRC4 repeat may promote filament formation via interactions involving the LFDE (Leu-Phe-Asp-Glu) region, with a binding that is not expected to perturb the inter-protomer interface. Conversely, the binding site for the FXXA (Phe-X-X-Ala) region is positioned at the RAD51 protomer-protomer interface and possesses filament-inhibitory potential (Short *et al*, 2016). Contact between the protein and the peptide is essentially held by hydrophobic interactions, with two phenylalanine residues playing a key role.

Nowadays, the only available parameter for molecular characterization of the BRCA2-RAD51 interaction is a rough estimation of the binding affinity obtained from a pull-down assay that claims a nanomolar affinity for this interaction (Jensen *et al*, 2010). Relative binding affinities have been reported for the interaction of RAD51 with each BRC repeat (i.e., BRC1-BRC8). Of the eight BRCA2 motifs, BRC4 shows the highest affinity for RAD51 (Cole *et al*, 2011). BRC4 has therefore always been used as a model to mimic the BRCA2-RAD51 interaction. Nevertheless, the only reported binding affinity of BRC4 for RAD51 is an indirect GST pull-down assay, which provides similar binding affinities for BRC1, BRC2, and BRC4 with an apparent K_*d*_ of 1-2 µM (Carreira & Kowalczykowski, 2011).

Most of the literature, comprising biophysical and structural characterizations, focuses on the interaction of BRCA2 with RAD51 fibrils formed on the DNA to unveil the details of the mechanism of DNA repair. Nonetheless, the BRCA2-RAD51 interaction, as a cancer-related target, can be disrupted directly in the cytosol at the RAD51 recruitment stage. If RAD51 self-assembled fibrils are not recruited as monomers by BRCA2, the HR activity is most likely inhibited due to the failure of RAD51 nuclear internalization. The intrinsic ability of RAD51 to form fibrillar structures (homo-oligomers) in vitro, also in the absence of DNA, has already been reported (Davies & Pellegrini, 2007; Holt *et al*, 2008). Nonetheless, these RAD51 fibrils recruited and monomerized by BRCA2 to be internalized in the nucleus as a response to DNA damage have been poorly characterized and understood.

Here, for the first time, we provide a comprehensive in vitro biophysical characterization of the native, self-assembled RAD51 fibrils and of their mechanism of disassembly in the presence of the BRC4 peptide (i.e., the fourth BRC repeat of the BRCA2 protein). BRC4’s mechanism of action is investigated in a concentration-dependent manner as the ability to disassemble RAD51 native fibrils utilizing complementary biophysical approaches. The detailed understanding of this mechanism is critical to design a specific inhibition of RAD51 recruitment at the site of DNA damage, leading to the HR failure in cancer cells that rely also on this pathway for survival. Then, based on cellular experiments, we propose BRC4 itself as a potential inhibitor of the HR pathway to improve cancer treatment. Indeed, the peptide is able to block HR in pancreatic cancer cells, hence providing a proof-of-concept of BRC4’s ability to trigger synthetic lethality (Kaelin, 2005) and anti-cancer activity.

## Results

The full-length recombinant human RAD51 protein was expressed in *E. coli* in a N-terminal 6xHis-tagged form and purified (see Materials and Methods). In its monomeric form, under reducing and denaturing conditions, it shows a molecular weight of about 40 kDa (Appendix Fig. S1). RAD51 though has a tendency to oligomerize. The size of its oligomers was first assessed via Static Light Scattering (SLS). The Zimm plot (Figure 1A) is the most popular method of SLS analysis. It allows one to estimate the radius of gyration (Rg), the second virial coefficient (A2), and the weight-average mass of the object under analysis. These data are reported in Table 1 for RAD51.

**Table 1.**
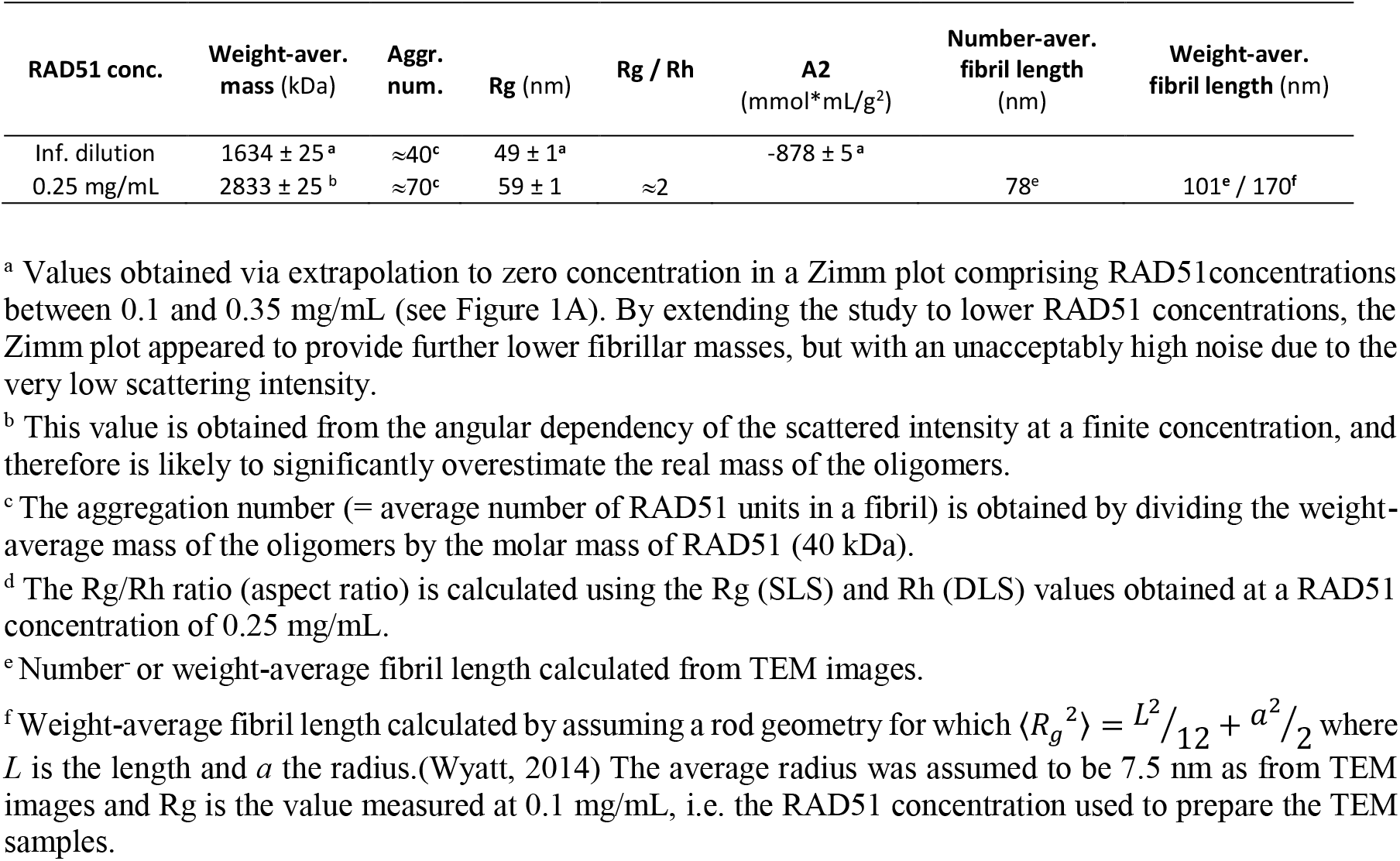
Structural features of RAD51 oligomers extrapolated at infinite dilution and at 0.25 mg/mL (6.25 µM).

**Figure 1.**
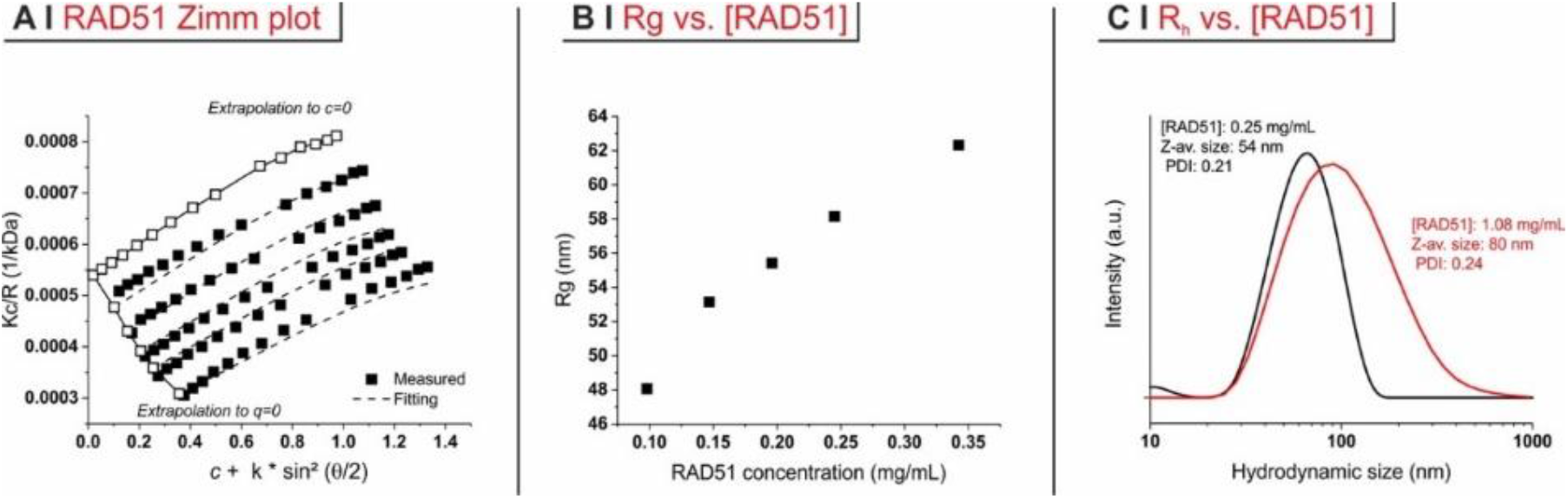
RAD51 fibrils characterization. ***A***. Zimm plot for RAD51 in a modified HEPES buffer at pH 8; *q* is the scattering vector (=4π/λ*sin(θ/2)). ***B***. Rg values derived from the angular dependency of scattering intensity are reported as a function of RAD51 concentration. ***C***. DLS intensity size distributions for RAD51 at 0.25 mg/mL (black) and 1.08 mg/mL (red).

The negative value of A2 indicates a high propensity to oligomerize. This is due to the RAD51 tendency to form fibrils. However, it suggests that the size of the oligomers may rather sharply depend on concentration. Indeed, while Rg value at infinite dilution is 49 nm – with fibrils comprising roughly 40 protomers each (Table 1) - ‘real’ solutions with [RAD51] ranging from 0.10 to 0.35 mg/mL show that Rg increases with concentration well above 60 nm (Figure 1B). A growth in size with increasing concentration can also be seen in Dynamic Light Scattering (DLS) measurements (Figure 1C,). From a morphological point of view, negative staining observations show that oligomers obtained at 0.10 mg/mL have a very elongated worm-like morphology (Figure 2A), which is confirmed by the high value of the aspect ratio as estimated through the Rg/Rh ratio (Table 1).

**Figure 2.**
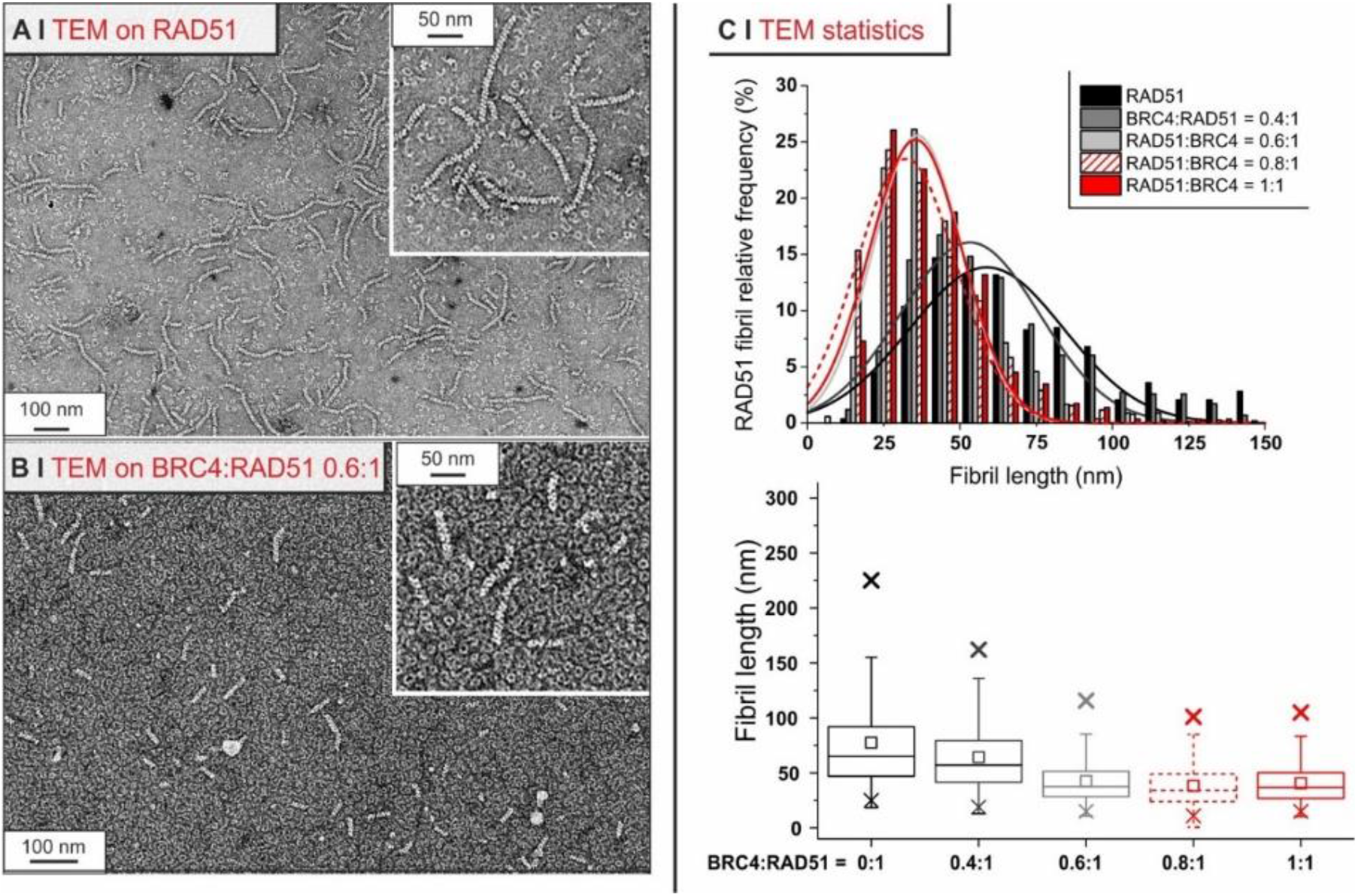
Negative staining TEM of RAD51 fibrils and their respective size statistics. ***A***. RAD51 recombinant protein as purified at a concentration of 0.1 mg/mL. ***B*** .RAD51 recombinant protein at a concentration of 0.1 mg/mL in the presence of BRC4 at RAD51/BRC4 1:0.6 molar ratio. ***C***. RAD51 fibril length number distributions with corresponding Gaussian curve fitting (up) and fibril length box plot (bottom) as a function of BRC4 concentration.

Fibril length estimates calculated from TEM images (black bars in Figure 2C) measurements suggest that the vast majority of the fibrils are likely to be 80-200 nm long.

To characterize BRC4’s effect on RAD51 fibrils, we exposed the fibrils to BRC4. Negative staining analysis showed a progressive decrease of fibril size with increasing BRC4 concentration, and fibrils eventually disappear when the ratio is significantly higher than the stoichiometric equivalence (Figure 2C). This was confirmed by size exclusion chromatography (SEC), which indicated that the material was mostly converted to small-size aggregates and monomers (Figure 3A), and in a more quantitative fashion by both SLS and DLS, which showed a clear decrease in fibril dimensions (namely Rg (SLS) and Rh (DLS)) above a BRC4:RAD51 molar ratio of 0.5 (Figure 3B).

**Figure 3.**
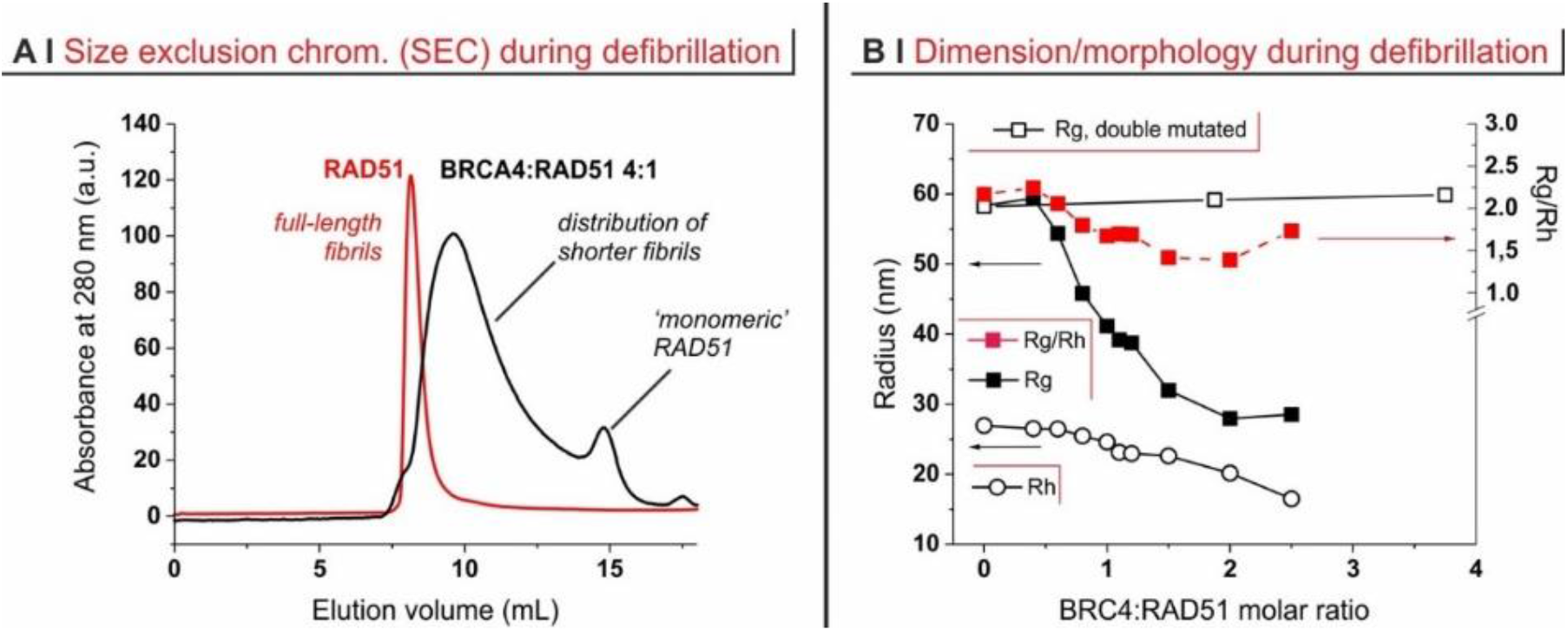
Characterization of BRC4’s effect on RAD51 fibrils. ***A***. Size exclusion chromatography (SEC) analysis of RAD51 before (*red line*) and after (*black line*) incubation with BRC4 (1 h, 37 °C; RAD51:BRC4 = 1:4). Please note that the ‘monomeric’ peak of RAD51 is caused by this form being the terminal step of the defibrillation process. ***B***. Rg, Rh and their ratio (aspect ratio) of RAD51 fibrils (0.25 mg/mL) upon addition of BRC4 or of scBRC4.

Notably, a double mutated version of BRC4 (scBRC4), where two amino acids critical for RAD51 binding (F1524, T1526) (Nomme *et al*, 2008; Scott *et al*, 2016) were replaced by alanine residues did not affect fibril size even with significant stoichiometric excesses of the mutated peptide (empty squares in Figure 3B). Additionally, TEM analysis showed a progressive decrease in RAD51 fibril size in the presence of increasing BRC4 concentration (Figure 2C).

Since BRC4 clearly shortens RAD51 fibrils, we further investigated this defibrillation phenomenon using techniques specifically sensitive to size/mass. In this context, SLS makes it possible to estimate the overall apparent weight average mass 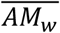 of a collection of objects in a colloidal dispersion. This is the linear combination of the masses of all objects in the dispersion, each weighted by their weight fractions (see Eq.1 Appendix Materials and Methods). In our case, the objects present in the colloidal dispersion are the RAD51 fibrils, the BRC4 peptide, and the BRC4-RAD51 complex. In the absence of the peptide, fibrils have a weight fraction of 1, which means that 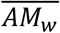 is the average mass of the fibrils. Upon addition of a compound unable to interact with the fibrils, 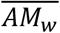 would slowly become smaller because the weight fraction of the fibrils (but not their size) would decrease. This effect is depicted by a dashed line in Figure 4A and confirmed with a BRC4 double mutant (empty squares in Figure 4A).

**Figure 4.**
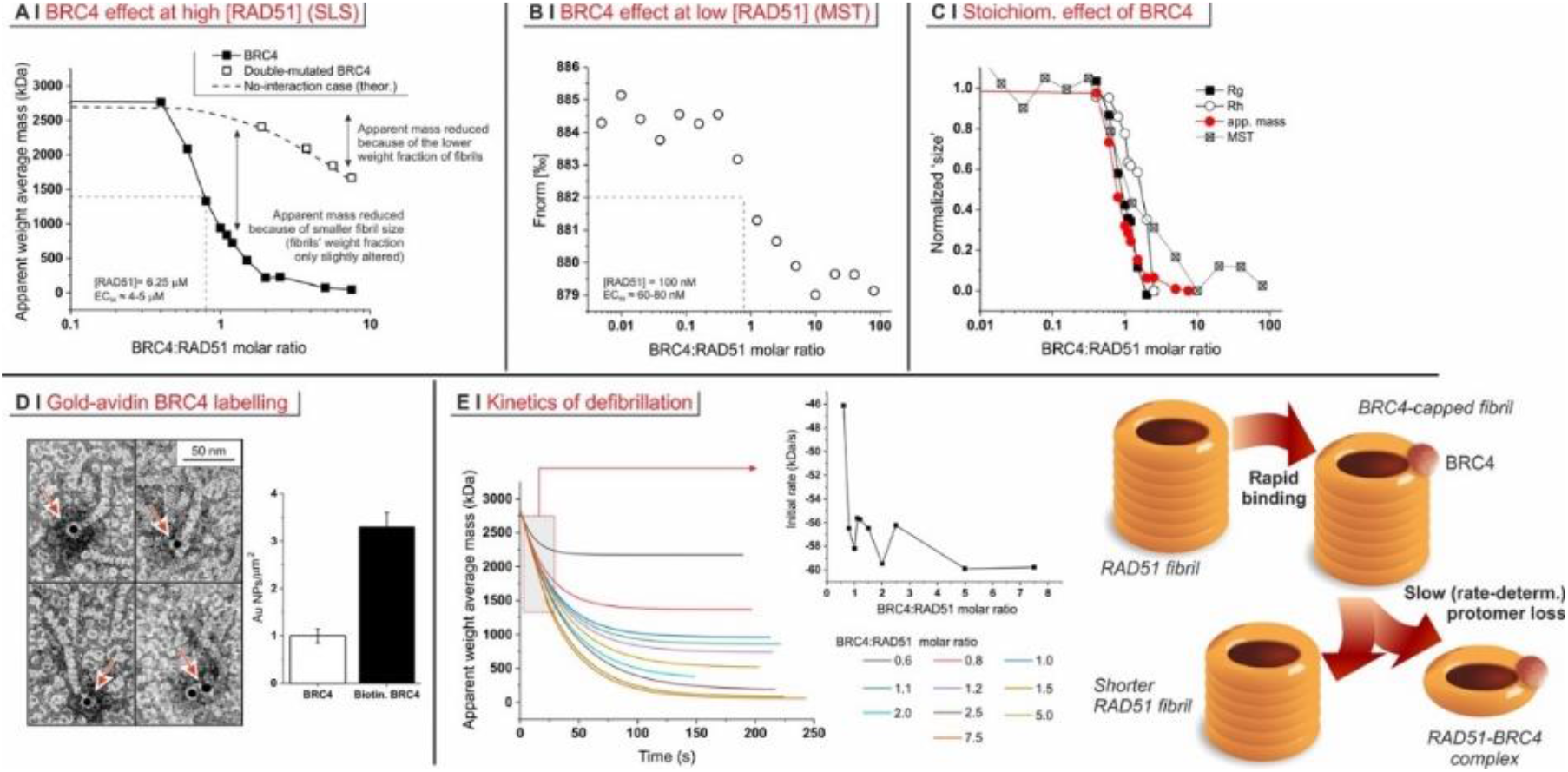
Mechanism of RAD51 defibrillation operated by BRC4. ***A***. Overall apparent weight average mass 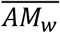 from SLS measurements on RAD51/BRC4 suspensions ([RAD51] = 6.25 µM = 0.25 mg/mL) as a function of the peptide/protein ratio. The dashed line is calculated assuming that a compound of the same mass of BRC4 is added and has no interaction with RAD51. The empty squares refer to SLS measurements performed with a double-mutated form of BRC4. ***B***. MST measurements performed on RAD51/BRC4 suspensions ([RAD51] = 100 nM = 4 µg/mL) as a function of the peptide/protein ratio. The data map closely with those of apparent mass 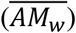, Rg and Rh (*inset*). ***C***. Localization of biotinylated BRC4 peptide by avidin-gold labelling. The histogram on the right shows the different densities of gold nanoparticles when non-labelled BRC4 or its biotinylated were used. The asterisk indicates a statistically significant difference (two-tailed Student t test *P<0.01). The labelled peptide is predominantly localized at only one of the fibril termini (23%, *n* = 67). In the remaining cases, it appears to be bound to isolated RAD51 monomers (73%, *n* = 67). ***D***. 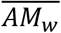 recorded as a function of time on RAD51/BRC4 suspensions ([RAD51] = 6.25 µM = 0.25 mg/mL) with various peptide/protein ratios. The data are fitted with an exponential function. The initial decrease rate, recorded as a function of the BRC4/RAD51 molar ratio (*inset*), is obtained as the derivative of the function at time zero. The pictorial sketch of the defibrillation mechanism considers a toroidal morphology of RAD51, as evidenced by TEM images in Figure 2.

Indeed, a non-interacting peptide decreases 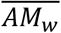 slowly and only when used in large amounts. In the conditions used for the SLS experiments ([RAD51] = 6.25 µM = 0.25 mg/mL), the midpoint of the process is close to the BRC4/RAD51 stoichiometric equivalence. This can largely be ascribed to the fibrils becoming shorter, and tallies nicely with the TEM observations. Since the details of RAD51 fibrillation (e.g. the fibril size) appear to be concentration-dependent, this relation may change at different [RAD51]. We thus explored the defibrillation at considerably higher dilution via microscale thermophoresis (MST). MST allows to quantify biomolecules interactions through thermophoretic detection of binding induced changes of a molecule (Jerabek-Willemsen *et al*, 2011). We used 100 nM (4 µg/mL) RAD51, previously labelled with a red fluorescent NT-647 dye specifically binding the N-terminal hexahistidine RAD51 tail (see Materials and Methods). Its titration with BRC4 (Figure 4B) showed again that the midpoint of the process was reached close to the BRC4/RAD51 stoichiometric equivalence.

Indeed, MST data closely mapped those from SLS (Rg and 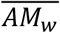) and DLS (Rh), all providing the same indication (Figure 4C). The relatively large amount of BRC4 needed to observe significant effects on fibrils may indicate a mechanism of defibrillation; whether this proceeds from a fibril terminus or is based on a random attack of BRC4 was not clear. We used TEM analysis of RAD51 fibrils in the presence of biotinylated BRC4 incubated with streptavidin-coated gold nanoparticles (10 nm) to further investigate the defibrillation mechanism (see Materials and Methods). More than a quarter of the nanoparticles appeared bound to fibrils (the rest to isolated RAD51 units, which are therefore to be considered RAD51/BRC4 complexes) and predominantly localized at only one of their termini (Figure 4D).

To further elucidate the mechanism of BRC4-induced defibrillation, we monitored its kinetics via SLS in experiments conducted at a constant RAD51 concentration (Figure 4E). Notably, the defibrillation rate appeared to be largely independent of the amount of BRC4. This may be because BRC4 binding is a rapid and likely equilibrium event, which means that BRC4-capped RAD51 fibrils may be relatively stable structures (hence they are visible through gold labelling in TEM). Their slow decomposition would therefore be the rate-determining event of the defibrillation process.

From a thermodynamic point of view, defibrillation is not a single equilibrium, but a succession of coupled steps (each with its own dissociation constant, *k*_*d*_). The whole process can be characterized by an EC_50_ value, i.e., the BRC4 concentration where the apparent fibril mass is halved, rather than a well-defined dissociation constant. It should be noted that the EC_50_ value depends on RAD51 concentration, due to the roughly stoichiometric nature of the effect; for example, with [RAD51] = 6.25 µM, the EC_50_ is about 4-5 µM (Figure 4A), while with [RAD51] = 100 nM the EC_50_ is 60-80 nM (Figure 4B).

We employed a mathematical model to extract some thermodynamic parameters from 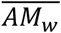 data. Of note, it is a semi-quantitative approach, due to the use of several important assumptions: i) independence of *k*_*d*_ on fibril length, i.e., we assumed all steps to have the same dissociation constant; ii) monodispersity of original fibrils, i.e., at the beginning of the process we ignore the fibril size distribution. Our simulation clearly showed a decrease in fibril length (number of protomers per fibril) with increasing BRC4/RAD51 molar ratio (Figure 5A); the products of intermediate defibrillation steps had a large dispersity in size, which is a common feature of all step-based processes, such as step-growth polymerization.

**Figure 5.**
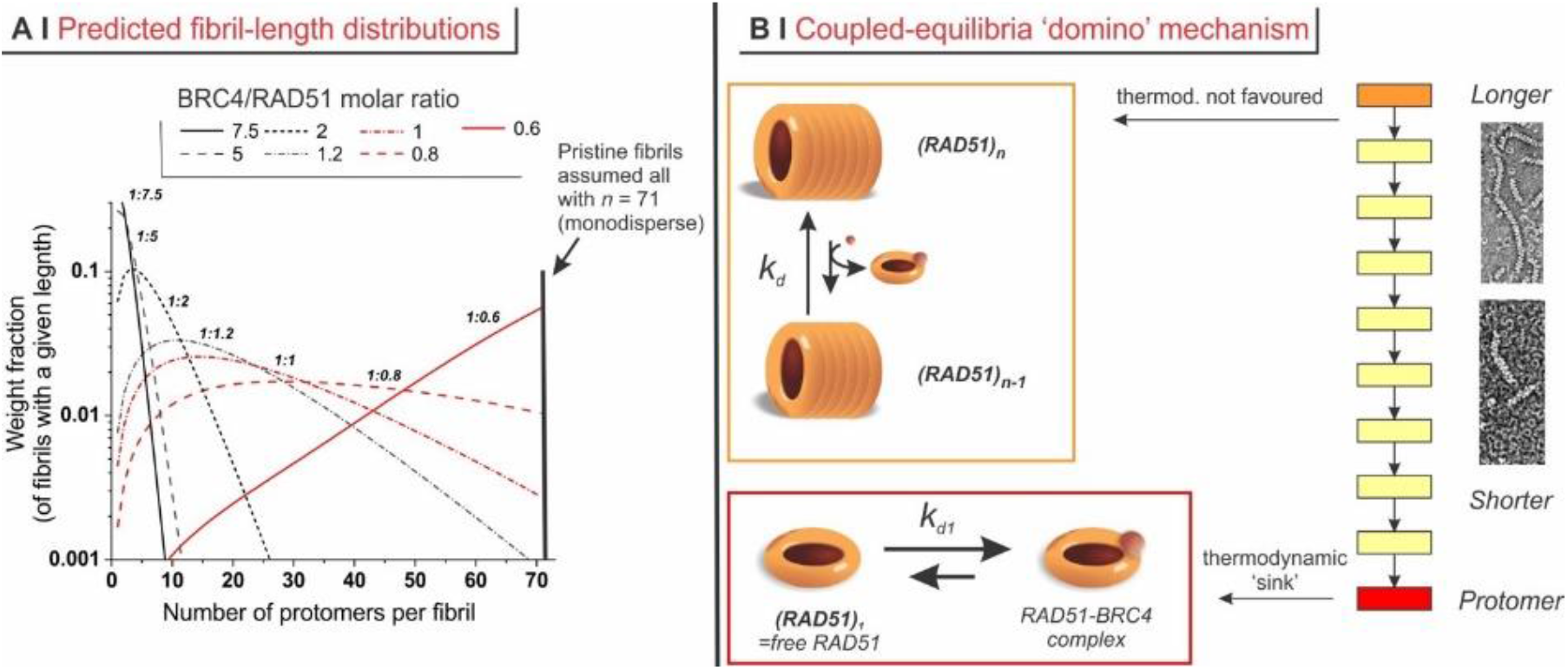
Simulation of the mechanism of RAD51 defibrillation operated by BRC4. ***A***. Simulated weight distribution of RAD51 fibrils with different lengths produced upon addition of variable amounts of BRC4. The model used is described in Appendix Materials and Methods and is fed with 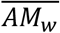 data. ***B***. Mechanistically, the whole process is based on a succession of losses of a RAD51 unit; each individual step appears to be characterized by a rather unfavorable constant. However, their coupling and above all the presence of a final ‘sink’ drive the process to completion: the complexation of RAD51 with BRC4 appears indeed to be thermodynamically favored, with a dissociation constant *kd1* calculated by our model to be mostly in the range 0.1-1 µM.

The *k*_*d*_ calculated for all BRC4/RAD51 molar ratios turned out to be essentially constant, with a value of 33.7 ± 0.5 (dimensionless, please see the section ‘Defibrillation model’ in Appendix Materials and Methods), which indicates that the individual steps - but the last - may not to be thermodynamically favored. Since the last event is the thermodynamically favored complexation of free protomer (released from its dimer), this may indeed act as the thermodynamic ‘sink’ that drives the whole chain of coupled steps (Figure 5B).

In the cytosol, RAD51 is thought to be present in a homo-oligomeric form. We have up to here used the BRC4 peptide to characterize in vitro the interaction between self-assembled RAD51 fibrils and BRCA2. According to the present data, BRC4 is able to efficiently bind RAD51 at the same site where BRCA2 binds and monomerizes RAD51. Since the peptide has no NLS, it most likely sequesters RAD51 in a monomeric form preventing its recruitment to the site of DNA damage. The data suggest that the BRC4 peptide may be a promising inhibitor of RAD51-mediated HR activity and as a consequence to prevent the repair of DNA damages that cancer cells frequently undergo.

BRC4’s ability to inhibit HR activity was evaluated in cells using an assay that relies on the recombination rate between two transfected plasmids (Abbott *et al*, 1998). A human pancreatic adenocarcinoma cell line that expresses fully functional BRCA2 (BxPC3) was selected as model for cell-based experiments due to its clinical relevance. In these cells, BRC4 seems unable to impair the HR pathway (Figure 6).

**Figure 6.**
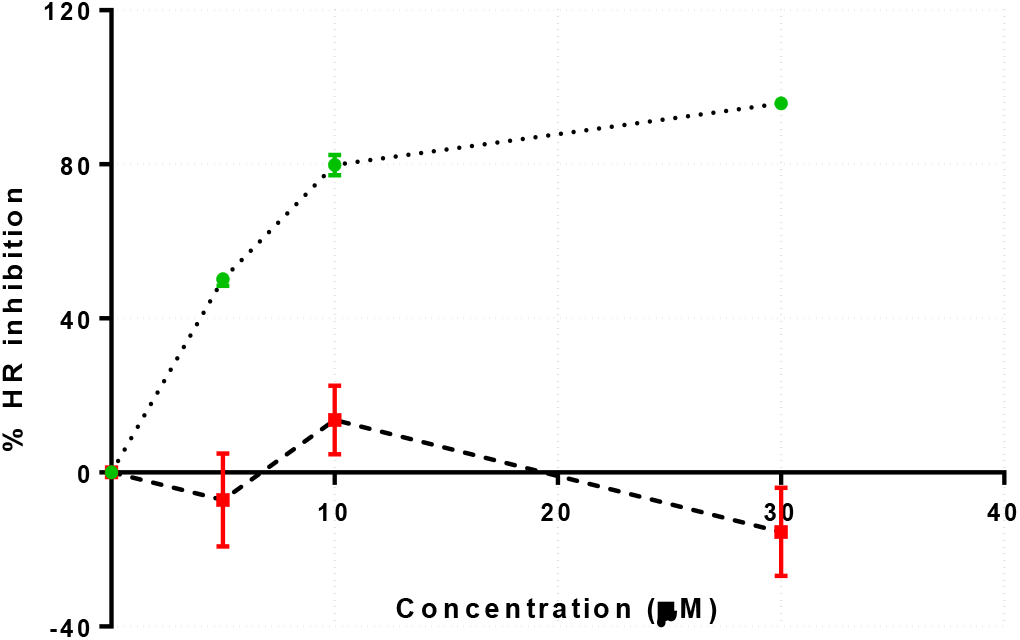
Homologous Recombination (HR) assay. Evaluation of the percentage of HR inhibition at 5, 10, and 30 μM BRC4 peptide (red) or BRC4 myristoylated peptide (green).

Nevertheless, the data could have been biased by a failure in cell membrane translocation of the peptide. To increase the intracellular release of the BRC4 peptide, a myristate group was attached to BRC4 (Nelson *et al*, 2007). A cysteine was added to the C-terminus of the BRC4 peptide and then conjugated to a myristoylated 5-amino-acid sequence (Cys-Lys(Myr)-Lys-Lys-Lys) via a disulfide bond formed between the cysteines of the two peptide domains. The disulfide bond of the resulting peptide (BRC4-ss-Myr) is cleavable (through reduction) upon entering the cell, thus freeing the cargo from its membrane tether (Nelson *et al*., 2007). Indeed, the advantage provided by the myristoylated peptide is that, once inside the cell, the myristoylated tag is cleaved upon disulfide bond reduction and the resulting peptide is in its native form with only one additional Cys at the C-terminal. Moreover, this intra cellular cleavage leads to an accumulation of BRC4 peptide in the cell that enhances the peptide inhibition activity, since once the myristoylated tag is cleaved the peptide cannot cross the cell membrane back.

Myristoylated BRC4 (BRC4myr) inhibited the HR pathway by up to 96.2% at 30 µM, with a clear dose-response curve (IC_50_ could be estimated around 5 µM), providing a proof of concept for the biophysical studies reported above. Moreover, we demonstrated the pivotal role of the myristoylation in providing BRC4 with the ability to cross the cell membrane (Figure 6).

Additional evidence of compromised HR in vivo was obtained by assessing the localization of RAD51 in BxPC-3 nuclei in response to DNA damage. To induce DNA damage, BXPC3 cells were treated with cisplatin (CPL 50 µM), which efficiently increased the number of cells bearing nuclear RAD51 foci (Figure 7A, B).

**Figure 7.**
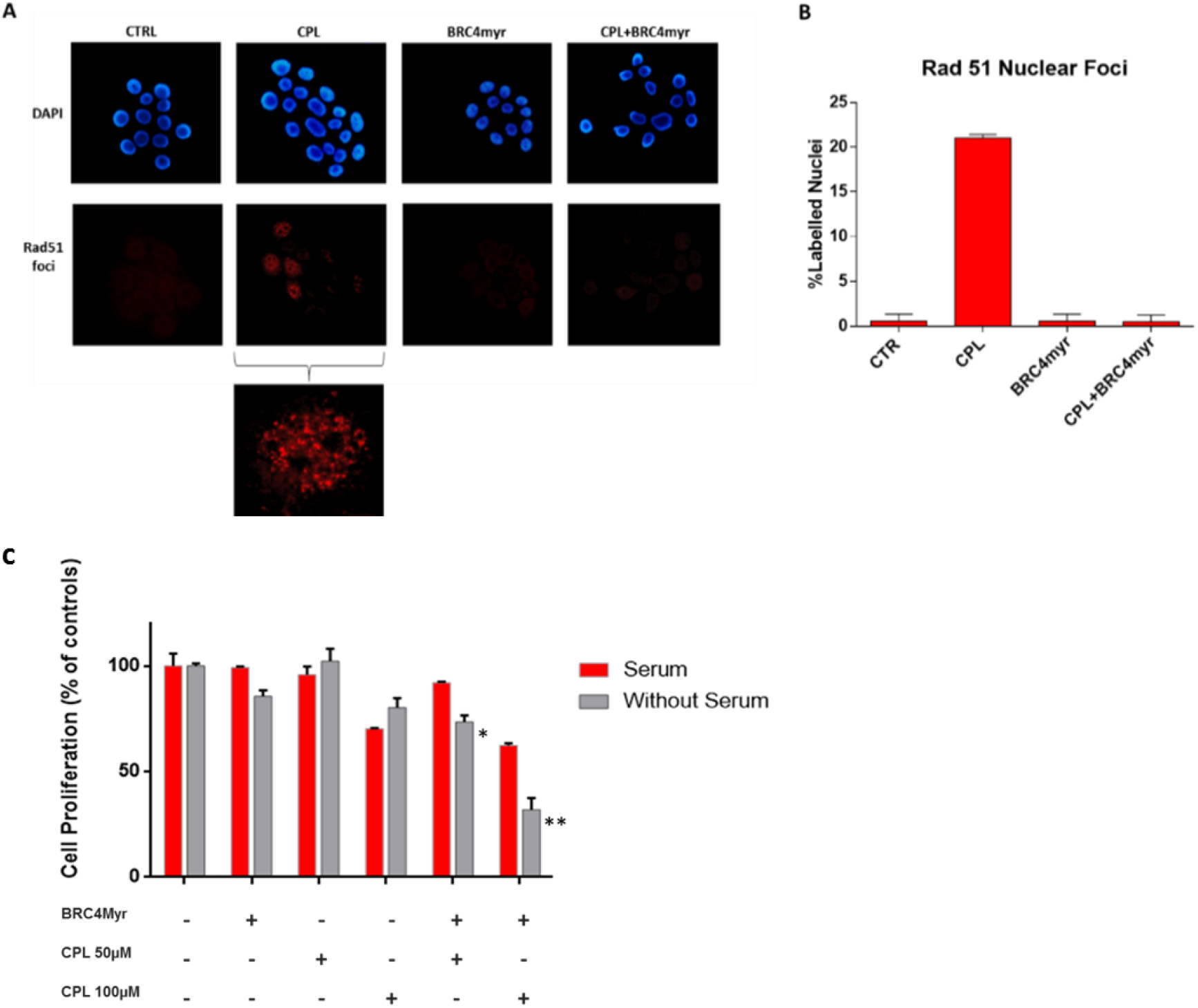
Immunofluorescence detection of RAD51. BxPC-3 cells were exposed to 50 μM cisplatin (CPL) given alone or in combination with 10 μM myristoylated BRC4 (BRC4myr). ***A***. Representative pictures showing DAPI stained cell nuclei and the corresponding immune-labeling of RAD51 localization. RAD51 labeling appears mainly in CPL-exposed cells and nuclear foci of RAD51 are clearly evident in 4 out of the 19 nuclei shown. A higher magnification detail of the CPL-treated sample is included. Foci are not detectable in untreated (CTRL), BRC4myr, and CPL+BRC4myr treated cells. ***B***. The bar graph shows the percentage of RAD51-labeled nuclei estimated by analyzing 300−400 cells for each treatment. Data were statistically evaluated by applying the one-way ANOVA, which indicated a significantly increased nuclear RAD51 labeling caused by CPL. ***C***. Cell viability assessed in BxPC-3 cultures exposed to CPL (50 or 100 µM, 1.5 h) and treated for 24 h with 5 µM BRC4myr. During the experiment, cultures were maintained in a medium with or without serum supplementation to ascertain a possible interference of serum with peptide effects. Data were analyzed by one-way ANOVA followed by Bonferroni post-test. In cultures maintained in the serum-deprived medium, BRC4myr significantly increased the effects of both CPL doses, with p < 0.01 (*) and < 0.001 (**), respectively.

The presence of BRC4myr strikingly suppresses formation of RAD51 foci in response to CPL treatment and restores the values of the nuclear foci to the untreated levels (CTRL) (Figure 7B). These data highlight the ability of the BRC4 peptide, in the presence of DNA damage, to inhibit RAD51 nuclear localization in cancer cells. The absence of RAD51-labeled nuclei detected in cultures exposed only to BRC4myr (10 μM) suggests that the peptide itself does not induce any detectable DNA damage.

We further validated the HR inhibitory activity of BRC4myr by investigating the effect induced by the peptide in cultured cells treated with CPL. After 1h pre-incubation with BRC4myr, BxPC3 cultures were exposed for 1.5 h to CPL administered at 50 µM or 100 µM and maintained for further 21 h in the presence of BRC4myr. The peptide concentration used (5 µM) is close to the IC_50_ dose assessed from the HR assay (Figure 6). Proteolytic degradation of peptide-based drugs is often considered a major weakness that limits their therapeutic applications (Bottger *et al*, 2017). Therefore, the experiment was performed using culture medium with or without serum supplementation. BRC4myr enhanced the antineoplastic effects produced by CPL in both doses (50 µM CPL, p<0,01; 100µM CPL, p<0,001), but this result was exclusively observed in the absence of serum (Figure 7C).

These data confirm the serum as a major factor hindering the effects of pharmacologically active peptides. However, in the absence of serum, the synergistic effect of the peptide with CPL is prominent. These findings strongly suggest that BRC4myr interferes with RAD51 nuclear localization and therefore HR repair, significantly affecting cell proliferation and leading to a synergistic effect with CPL. The above results support the proposed mechanism of action for BRC4 inhibition.

## Discussion

We have here shown that recombinant RAD51, in the absence of DNA, form worm-like fibrils that are mostly 100-200 nm in size, notwithstanding these dimensions depend on its concentration. It seems reasonable to hypothesize that this dependency is caused by an equilibrium of terminal association between RAD51 protomers and fibril termini. We are inclined to exclude the idea that higher concentration increases nucleation of new fibrils. This is because it would cause a fraction of smaller fibrils at higher RAD51 concentration (as happens for fibrin (Leon-Valdivieso *et al*, 2018)).

Here, we have ample indications that BRC4 binds to RAD51 and that this binding reverts the fibrillar self-association of RAD51

Mechanistically, our results suggest that BRC4 may disassemble fibrils gradually from their termini rather than through attacks at random positions along the fibril. In fact, all indicators of fibril size (mass, dimension, diffusion coefficient) decreased very gradually, with the midpoint of the process being observed close to the 1/1 BRC4/RAD51stoichiometry. Further, TEM, SLS, and DLS have shown that the fibrils preserved an elongated morphology (a high-aspect ratio) during the process, and BRC4 localized at the fibril termini. Finally, if defibrillation proceeds from fibril termini, the number of fibrils (and of their terminal groups) is constant throughout the process. Importantly, this may not be the only mechanism of interaction between RAD51 and BRC4, although possibly the most important at high peptide concentration: at sub-stoichiometric amounts, BRC4 has been reported to promote RAD51 fibril nucleation (Liu *et al*, 2010), which may indicate two modes of interaction. In one, BRC4 would bind the core catalytic domain of (multiple) RAD51 protomers, facilitating their assembly, while in the other BRC4 would bind at the site of protomer-protomer interactions, disrupting them (Short *et al*., 2016).

Kinetically, a 2-stage mechanism appears likely: the rate of defibrillation does not depend on BRC4 concentration, which may be explained with a rapid BRC4 terminal binding possibly followed by a rate-determining loss of a BRC4-bound protomer unit. A similar mechanism was previously suggested for the assembly of RAD51 fibrils on DNA assisted by BRCA2 (Nomme *et al*., 2008; Shahid *et al*., 2014). From a thermodynamic point of view, the rather high value of *k*_*d*_ calculated for the individual steps of RAD51 loss indicates that the driving force of the process may not be a high affinity of the peptide for its fibrillar substrate, but rather a ‘domino’ combination based on i) the coupling between successive RAD51 losses, so that the removal of RAD51 from an *n*-mer shifts to the right all equilibria involving larger fibrils; ii) the thermodynamically favored complexation of BRC4 to free protomer, which acts as a final ‘sink’.

The demonstration that BRC4 binds to RAD51 with dramatic (but close to stoichiometric) morphological consequences is key to interpreting the biological finding that the peptide inhibits the HR pathway once internalized into the cell. Indeed, BRC4 can bind RAD51, but cannot further support RAD51 activity within the HR pathway as is physiologically done by BRCA2 (Holt *et al*., 2008) because it lacks the BRCA2 NLS. In fact, even upon DNA damage (CPL), RAD51 is not able to localize at the site of damage in the presence of BRC4. This is in contrast to the RAD51 foci formation observed when the peptide is not present. BRC4 can compete with BRCA2 for RAD51 binding, being however unable to support the HR pathway as BRC4 lacks NLS. It can act as an inhibitor and arrest the DNA repair mechanism. Indeed, it was able to potentiate the CPL effect, inhibiting the mechanism of DNA repair and more efficiently inhibiting proliferation of pancreatic cancer cells.

A recent study has reported a cell-penetrating peptide (CPP) comprising a 16-amino-acid stretch of the BRC4 repeat, which can inhibit RAD51-BRCA2 interaction (Trenner *et al*, 2018). This peptide prevents RAD51 loading onto single-stranded DNA, and thus causes defective homology-mediated repair of DSBs and increased degradation of stalled DNA replication forks. However, the affinity of this peptide for RAD51 has not yet been reported. Affinity data have only been reported for small tetrameric sequences derived from the BRC4 repeat (Scott *et al*, 2013). In particular, the affinity of the FHTA peptide (native BRC4 sequence) for RAD51 is in the low millimolar range, which is consistent with a fragment peptide that has to be evolved towards a full efficient peptide inhibitor.

Small organic molecules mimicking BRC4 and inhibiting RAD51-BRCA2 interaction have also been reported in the literature (Bagnolini *et al*, 2020; Falchi *et al*., 2017; Roberti *et al*, 2019; Zhu *et al*, 2013). The molecular mechanism here reported can eventually shed further light on the complex HR pathway, which inhibition, through peptides or small molecules, could be rather key for the discovery of innovative anticancer compounds.

Here, we have characterized the BRC4 peptide as an effective inhibitor of the BRCA2-RAD51 binding with a significant effect on RAD51 fibril disassembly and protein translocation. Nevertheless, the BRC4 peptide is not able to enter the cell. In the present study, we used peptide myristoylation to circumvent cell membrane internalization in vitro. However, this solution is not physiologically applicable due to proteolytic degradation. Nevertheless, myristoylation is a critical tool to internalize the peptide in the cell, and upon physiological cleavage, no potentially interfering tag is attached to the peptide.

The BRC4 peptide is therefore an excellent tool to characterize the disassembly process and a promising inhibitor of the BRCA2-RAD51 protein-protein interaction. In fact, we have proved its ability in cancer cells to strikingly prevent RAD51 nuclear localization upon DNA damage, consequently affecting cell proliferation after treatment with cisplatin. Based on these results, we now aim to improve the potency of the peptide and its drug-like properties. In particular, we plan to pursue smart formulation, nanoparticles, and intelligent deliveries to improve BRC4 drug-likeness.

## Materials and Methods

Complete experimental descriptions are reported in Appendix Materials and Methods.

### Protein expression and purification

The full-length human RAD51, with a 6xHis tag at the N-terminus, was expressed in *E. coli* Rosetta2(DE3)pLysS cells carrying a pET15b plasmid. The protein. *E. coli* cells were grown into a TB-5052 auto-induction medium at 20 °C for 72 h. The protein was purified according to the protocol reported in Appendix Materials and Methods.

### Physico-chemical characterization

#### Size exclusion chromatography (SEC)

The recombinant pure human RAD51 protein was incubated at 37 °C for 1 h in the absence or presence of the BRC4 peptide in a 1:4 molar ratio. After incubation, protein samples were loaded onto a size exclusion chromatographic column (Superdex200 Increase 10/300 GL, GE Healthcare).

#### Static light scattering (SLS) and composition-gradient multi-angle static light scattering (CG-MALS)

Measurements were carried out using a Calypso automated delivery system as a mixing unit, connected to a Dawn Heleos II multi-angle light scattering (MALS) and a T-rEX refractive index (RI) detector, both operating at 660 nm (all instruments produced by Wyatt Technology, Santa Barbara, California).

#### Dynamic Light Scattering (DLS)

100 μL of the RAD51/peptide mixtures were manually collected at the end of the analysis above and analyzed at a temperature of 25 °C with a Möbiuζ instrument (Wyatt Technology, Santa Barbara, California), which was equipped with a laser at 532 nm and operated at a scattering angle of 163.5°.

#### Microscale Thermophoresis (MST)

6xHis-RAD51 was labelled with the Monolith His-Tag Labeling Kit RED-tris-NTA 2nd Generation kit (NanoTemper Technologies), which specifically recognizes the hexahistidine tail of the recombinant protein.

#### Transmission Electron Microscopy (TEM)

A) Negative staining. Recombinant RAD51 protein (0.1 mg/mL) alone or in the presence of different concentrations of the BRC4 peptide were incubated for 2 h at 4 °C. After incubation, each sample was adsorbed into pure carbon film 300-mesh copper grids and negatively stained using 1% uranyl acetate (JEM-1011 (JEOL) TEM). B) Negative staining coupled with streptavidin-gold labeling. 5 μL drops of recombinant 6xHis-RAD51 protein (0.1 mg/mL) alone or in the presence of different concentrations of BRC4 were incubated for 30-90 s on plasma-cleaned carbon-film-coated 300 mesh nickel grids. After washing, grids were incubated for 30 min (at room temperature) or 2 h (at 4 °C) with streptavidin-gold diluted 1:20-1:60 in washing buffer. The grids were then washed, negatively stained, and imaged as described above.

### Defibrillation model

To improve our understanding of the disassembly mechanism of RAD51 fibrils, based on the obtained experimental data, we have employed a mathematical approach conceptually similar to those often employed in step-growth (equilibrium) polymerization. This is based on the estimation of dissociation constants through its kinetic equations, allowing the system to reach the steady state at each defibrillation step.

### Cell culture

BxPC-3 were grown in RPMI 1640 containing 10% FBS, 100 U/mL penicillin/streptomycin, 2 mM glutamine.

### Homologous Recombination assay

Homologous recombination (HR) was assessed using a commercially available assay (Norgen).

### Immunofluorescence

Immunofluorescence was used for studying RAD51 nuclear translocation. To visualize RAD51 in cell nuclei, BxPC-3 cells were seeded on glass coverslips placed in a 6-well culture plate (2 × 10^5^ cells/well) and allowed to adhere overnight.

### Cell viability assay

Cell viability was assessed by CellTiter-Glo Luminescent Cell Viability Assay from Promega.

## Acknowledgments

This work was supported by the Italian Association for Cancer Research (AIRC) through Grant IG 2018 Id.21386 and the Istituto Italiano di Tecnologia (IIT).

## Authors contribution

AC conceived the project with input from SG, NT, GDS and MR. FS, MM, RM, GDS and SG designed all in vitro and in cells experiments. FS performed and analyzed all in vitro experiments with the support of FR and with the following exceptions: AG performed and analyzed SLS and DLS experiments; RM performed and analyzed TEM experiments; AA performed the analytical characterization of purified proteins and peptides; MM performed and analyzed all in cells experiments. WR designed the defibrillation model. SG supervised project activities. FS, SG, AC, and NT wrote the manuscript.

## Conflict of interest

The authors declare that they have no conflict of interest.

## Supporting Information

Appendix (PDF document)

